# Remodeling and compaction of the inactive X is regulated by *Xist* during female B cell activation

**DOI:** 10.1101/2022.10.19.512821

**Authors:** Isabel Sierra, Son C. Nguyen, R. Jordan Barnett, Ashley L. Cook, Han-Seul Ryu, Zachary T. Beethem, Jennifer E. Philips-Cremins, Eric F. Joyce, Montserrat C. Anguera

## Abstract

X Chromosome Inactivation (XCI) equalizes X-linked gene expression between sexes. B cells exhibit unusually dynamic XCI, as Xist RNA/heterochromatic marks on the inactive X (Xi) are absent in naïve B cells, but return following mitogenic stimulation. Xi gene expression analysis supports dosage compensation, but reveals high levels of XCI escape genes in both naive and activated B cells. Allele-specific OligoPaints indicate similar Xi and Xa territories in B cells that is less compact than in fibroblasts. Allele-specific Hi-C maps reveal a lack of TAD-like structures on the Xi of naïve B cells, and alterations in TADs and stronger TAD boundaries at Xi escape genes after mitogenic stimulation. Notably, *Xist* deletion in B cells reduces Xi compaction and changes TAD boundaries, independent of its localization to the Xi. Our findings provide the first evidence that Xi compaction/small scale organization in lymphocytes impact XCI maintenance and female biased X-linked gene expression.

## INTRODUCTION

X Chromosome Inactivation (XCI) equalizes X-linked gene expression between the sexes in female placental mammals. XCI occurs during early embryonic development through a multistep process initiated by the expression of the long non-coding RNA Xist^1–4^. The choice of X for silencing in XCI is random, but the future inactive X chromosome (Xi) can be identified by upregulation and spread of Xist RNA transcripts and heterochromatic factors that remove active histone modifications along its length. Various repressive epigenetic modifications are then added to the Xi, resulting in gene repression across most of the Xi^5–11^. The enrichment of Xist RNA and heterochromatic modifications across the Xi maintain a memory of transcriptional silencing with each cell division that is maintained into adulthood. While most of the Xi is transcriptionally silenced, some genes on the Xi ‘escape’ XCI in female somatic cells^12, 13^. XCI escape genes usually lack Xist RNA and heterochromatic modifications^14–18^, supporting a critical role for Xist RNA and heterochromatic modifications to reinforce transcriptional silencing. Indeed, *Xist* deletion in a variety of somatic cells results in partial reactivation of the Xi, with cell-type specific abnormal overexpression of X-linked genes^19–22^. While recent studies have identified XCI escape genes across various mouse and human tissues^23, 24^, escape genes responsible for sex biased function, particularly in immune cells^25, 26^, that could contribute to female-biased autoimmune diseases, are not well defined.

While the active X (Xa) retains typical features of mammalian chromosomes, including A (open/active chromatin) and B (closed/repressed chromatin) compartments, Hi-C generated allele-specific spatial proximity profiles from mammalian fibroblasts and neural progenitor cells (NPCs) suggested that the Xi lacks compartments and TADs^27, 28^. However, the presence of TADs on the Xi remains unresolved, as higher resolution Hi-C sequencing revealed faint, low-resolution TADs^29, 30^. In contrast to the Xa, the Xi is highly compacted and spherical^31^ and has a unique bipartite organization involving two ‘mega-domains’ separated by the microsatellite repeat *Dxz4* locus^27, 28, 32^. The unique three-dimensional structure of the Xi is thought to be dependent on Xist RNA, as *Xist* deletion in fibroblasts and NPCs results in loss of the mega-domain partition and appearance of well-defined TADs across the Xi^27^. While it appears that Xist RNA is required for maintenance of the three-dimensional structure of the Xi, the mechanistic basis for this effect remains unclear.

Most somatic cells have persistent enrichment of Xist RNA and heterochromatic modifications at the Xi that are cytologically visible using RNA fluorescent in situ hybridization (FISH) and immunofluorescence (IF) techniques^33, 34^. However, female lymphocytes exhibit an unusual dynamic form of XCI maintenance, as canonical robust Xist/XIST RNA ‘clouds’ are absent at the Xi of naive B cells from both mouse and human^35–37^, but Xist RNA re-localizes to the Xi following *in vitro* B cell stimulation with either CpG oligodeoxynucleotides (CpG) or lipopolysaccharide^36, 38^. Moreover, heterochromatic histone modifications H3K27me3 and H2AK119Ub missing from the Xi in naive B cells appear concurrently with Xist RNA following *in vitro* B cell stimulation^36, 38^. As *Xist/XIST* is constitutively expressed in naïve B cells, an inability to localize Xist RNA to the Xi in naïve B cells that can be overcome by B cell activation may provide a mechanistic explanation for the dynamic XCI maintenance observed in lymphocytes^36, 38, 39^.

Naive B cells are quiescent, with significantly reduced transcription rates and highly condensed chromatin^40^. However, whether transcriptional repression is preserved across the Xi in naive B cells lacking enrichment of Xist RNA and heterochromatic histone modifications is unknown, and the frequency of XCI escape genes has not yet been evaluated. Moreover, while *in vitro* B stimulation causes significant genome-wide chromatin decondensation and rearrangement within the nucleus, with increased short-range DNA contacts and looping, all before the first cell division^41, 42^, how the accumulation of Xist RNA impacts DNA contacts at silenced and XCI escape genes across the Xi in activated B cells is unknown.

Here we examine the transcriptional activity of the Xi in naive and *in vitro* activated female mouse B cells, to determine how activation impacts gene expression and the nuclear structure of the Xi. We find the Xi is dosage compensated despite the absence of Xist RNA accumulation across the Xi in naive B cells, and that 17-24% of expressed X-linked genes escape XCI in both naive and stimulated B cells. Using allele-specific OligoPaints and Hi-C analyses, we see that the global territory and compartmentalization of the Xi is relatively unchanged with B cell stimulation, yet we observe both dynamic changes in gene expression and Xi structure at the TAD level. Importantly, we show that likely both *Xist* transcription and Xist RNA transcripts are necessary for limiting stimulation induced changes to compaction of the Xi territory and Xi TAD organization in B cells. Together, these findings provide the first evidence that Xi compaction/small scale organization in lymphocytes may influence XCI maintenance and female biased X-linked gene expression, providing a potential mechanistic explanation for the sex bias observed in many autoimmune diseases where B cells are pathogenic.

## RESULTS

### The Xi is dosage compensated in the absence of Xist RNA localization in naive B cells

To determine if Xi dosage compensation is impacted by the lack of Xist RNA ‘clouds’ at the Xi, we compared transcription from the Xa and Xi in naïve and *in vitro* stimulated female B cells in which Xist RNA ‘clouds’ are restored at the Xi. For these studies, we used a mouse model of skewed XCI, in which a female *Mus musculus* mouse harboring a heterozygous *Xist* deletion is mated to a wild-type *Mus castaneus* male, generating F1 mice in which the paternal X chromosome is uniformly inactivated and single nucleotide polymorphisms (SNPs) can be used to distinguish each allele **(Figure 1A - left)**. Using these F1 mice, we prepared RNA from splenic CD23+ naive B cells and B cells stimulated *in vitro* with CpG for 24hr for allele-specific RNA sequencing **(Figure 1A – right)**. Principle component analysis of naive and *in vitro* stimulated B cells revealed separation between naive and stimulated samples, with tight clustering of replicates within each state **(Supplementary Figure S1A)**. Surprisingly, we observed that the Xi is largely dosage compensated in both naive and *in vitro* stimulated B cells, as exhibited by the significantly higher mapped reads (reads per million mapped (RPM)) from the Xa compared to the Xi **(Figure 1B)**. Xi gene expression arose from various regions across the Xi in both naive and in *vitro* stimulated B cells, with higher expression of genes located at the distal ends of the chromosome compared to the center **(Figure 1C)**. Expressed regions were similar between the Xa and Xi, with higher expression for the Xa, in both naive and *in vitro* stimulated B cells **(Supplementary Figure S1B)**.

**Figure 1.**
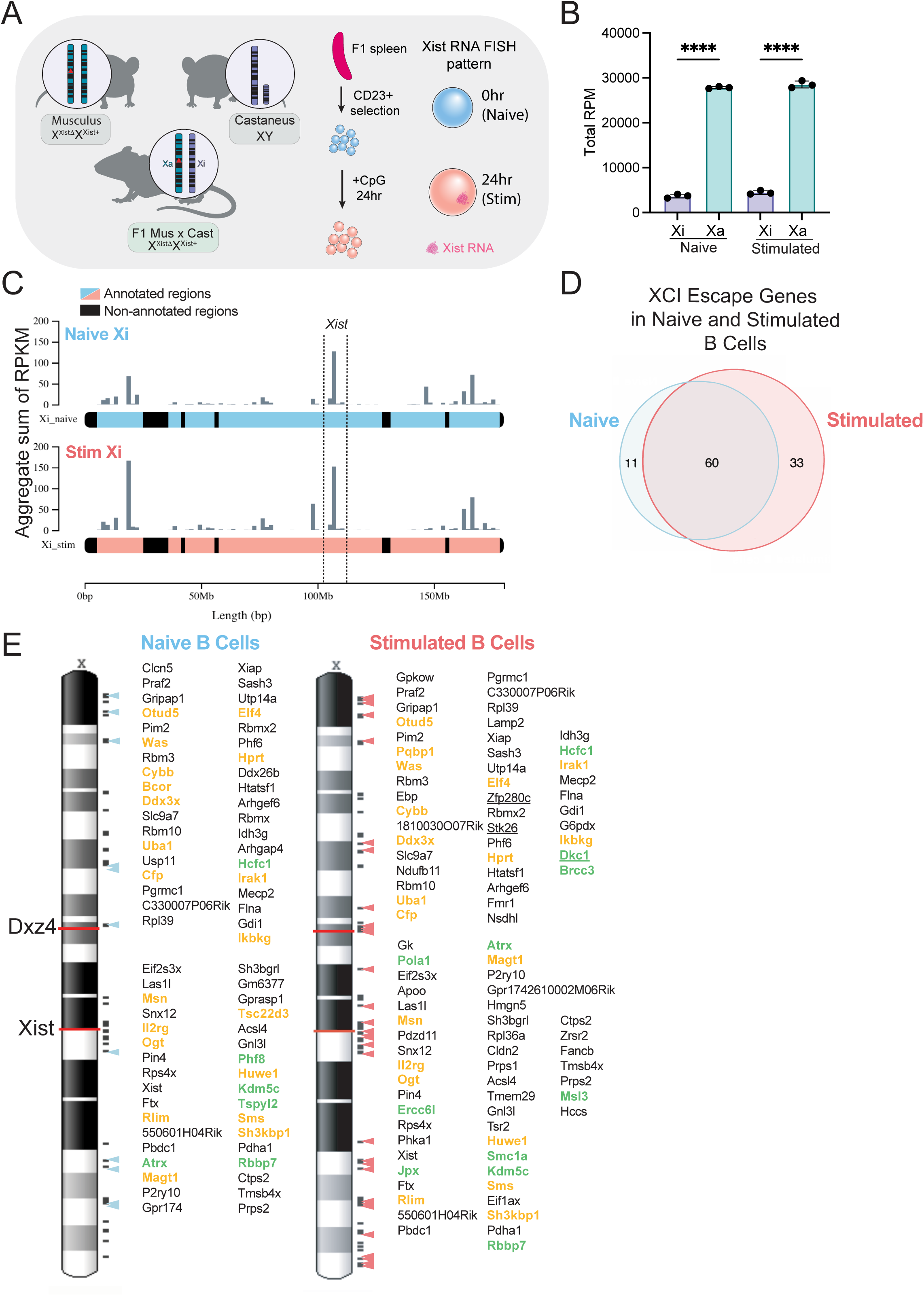
The inactive X in B cells is dosage compensated and has XCI escape genes in both naive and *in vitro* stimulated B cells. **A)** <u>Left</u> - Schematic for the F1 Mus x Cast mouse with completely skewed XCI. Female *Mus musculus* (Musculus) heterozygous for *Xist* deletion (Xistι1) is mated to a wild-type male *Mus castaneus* (Castaneus) mouse. The F1 generation expresses *Xist* exclusively from the Castaneus allele, thus the wildtype Xi is paternally inherited (Castaneus) in every cell of this mouse. <u>Right</u> – Schematic for the splenic follicular B cell isolation procedure used throughout this study. Naive CD23+ follicular B cells are stimulated *in vitro* with CpG for 24 hours. **B)** Allele-specific RNAseq analyses showing reads per million mapped (RPM) of X-linked reads mapping to either the Xi (Cast, purple) or Xa (Mus, teal) genomes in both naive and stimulated B cells. Bars represent mean +/- SD. Statistical test performed using One-way ANOVA with Tukey’s correction for multiple comparisons. ** p-value < 0.005, **** p-value < 0.0001. **C)** X chromosome plots generated by chromoMap^61^ showing the genomic location of Xi-specific expression in naive (blue) and stimulated (coral) B Cells. Values are displayed as the sum of RPKM in 1.7Mb bins. Dotted lines display location of the *Xist* locus. Colored regions represent annotated regions of the X chromosome, black bars represent un-mappable regions. **D)** Venn diagram showing the distribution of the 104 XCI escape genes identified in naive and stimulated B cells (11 naive specific; 33 stimulated specific; 60 shared). **E)** X chromosome maps showing the location of each XCI escape gene. Genes are listed in linear order as they appear across the X chromosome. Colored arrows indicate location of XCI escape genes for either naive (left) or stimulated (right) B cells. *Xist* and Dxz4 boundary are indicated with red lines. Genes colored in orange have immune-related functions. Genes colored in green have chromatin organization functions. Underlined genes are novel escapees identified in this study.

### XCI escape genes are present in naive and stimulated B cells

Using the allele-specific RNAseq datasets, we applied a previously published binomial distribution approach to identify XCI escape genes in female mouse B cells (see methods)^23^. Genes were considered expressed if their diploid Reads Per Kilobase Million (RPKM) was greater than 1 and their haploid expression (SRPM) was greater than 2. A 95% confidence interval was applied and an expressed gene was considered to have ‘escaped’ if the probability of escape was greater than 0. We found a total of 249 X-linked genes expressed in both naive and stimulated B cells, of which 104 genes (roughly 41%) could be considered XCI escape genes across both naive and stimulated B cells **(Supplementary Table 1)**. XCI escape genes were largely shared between naive and stimulated B cells (60 genes) and exhibited diverse expression patterns in both states (**Supplementary Table 1, Supplementary Figure S1C**). We identified 11 XCI escape genes unique to naive B cells, and 33 genes unique to stimulated B cells **(Figure 1D).** Unique XCI escape genes spanned the X, residing near clusters of other escape genes (colored arrowheads in **Figure 1E**). Using gene ontology analyses, we found that B cell specific XCI escape genes function in XCI regulatory pathways, RNA processing, nucleotide metabolism, ribonucleoprotein complex biogenesis, negative regulation of cellular component organization, regulation of cytoplasmic transport, and regulation of type I interferon production (**Supplementary Figure S1D**). Some novel XCI escape genes were specific for mouse B cells: *Dkc1, Zfp280c, Stk26,* (**Figure 1E**, underlined). There were 20 immunity-related XCI escape genes across naive and stimulated B cells, including *Was, Il2rg, Irak1, Cfp*, *Bcor*, and *Ikbkg* (**Figure 1E; genes in orange**). XCI escape genes also appear to function in transcription, chromatin modification, and regulation of chromosome architecture, including *Kdm5c, Smc1a, Jpx, Pol1a, Msl3, Phf8*, and *Atrx* (**Figure 1E**, genes in green). Intriguingly, a number of genes that regulate nuclear organization and chromatin architecture exclusively escape XCI in stimulated B cells (*Jpx, Smc1a, Ercc6l, Brcc3, Dkc1, Msl3 and Pola1*), suggesting a novel role for increased dosage of X-linked genes in genome-wide nuclear organization changes in female B cells following stimulation.

### Xist RNA transcripts are detected uniformly across the Xi at 12 hrs post-stimulation, and are enriched at the Xist locus at 24 hrs in stimulated B cells

To directly assess Xist RNA enrichment across the Xi, we performed Capture Hybridization of RNA Targets (CHART) in primary female naïve and *in vitro* stimulated B cells at 12 and 24 hours after CpG treatment. As expected, Xist RNA was not detectable across the Xi in naive B cells **(Figure 2B, 0 hr),** with levels identical to those for chromosomes 4 and 13, which lack Xist localization (**Supplementary Figure S2**). However, at 12 hours post-stimulation, Xist RNA transcripts were detected across the Xi, and enrichment increased uniformly at all these regions by 24 hours (**Figure 2B**). We quantified the Xist RNA enrichment levels at 12 and 24 hours post-stimulation at three regions: the Xist RNA locus itself, the region +/- 2Mb surrounding *Xist (*excluding the *Xist* promoter), and across the Xi. Xist RNA transcript accumulated at the *Xist* gene at 12 and 24 hours, with lower yet evenly distributed Xist RNA reads +/- 2Mb surrounding the Xist gene (**Figure 2B, inset**). Relatively similar levels of Xist RNA were observed across the Xi and around the *Xist* gene at 12 hrs (**Figure 2C**), suggesting that Xist RNA is uniformly tethered across the Xi, in contrast to what is observed during XCI initiation^43^. By 24 hours significantly more Xist RNA transcripts were present at the *Xist* locus compared to +/-2Mb and the entire Xi (**Figure 2C**), and predominately accumulated at distal intergenic regions (46.80%) and intronic regions (13% at first intron, 28.52% at other introns) **(Figure 2D)**, similar to what is observed in differentiated embryonic stem cells and mouse embryonic fibroblasts^15^. We also found eight regions across the Xi that were significantly depleted for Xist RNA transcripts **(Figure 2A, blue vertical bars)** at both 12 and 24 hours post-stimulation (**Figure 2E**). These regions included 92 X-linked genes (**Supplementary Table 2**), about a third (31 genes) of which were XCI escape genes **(Supplementary Table 2– green highlights)**. The remaining 71 genes were transcriptionally repressed, but lacked Xist RNA enrichment, suggesting Xist RNA independent mechanisms for gene silencing at these loci.

**Figure 2.**
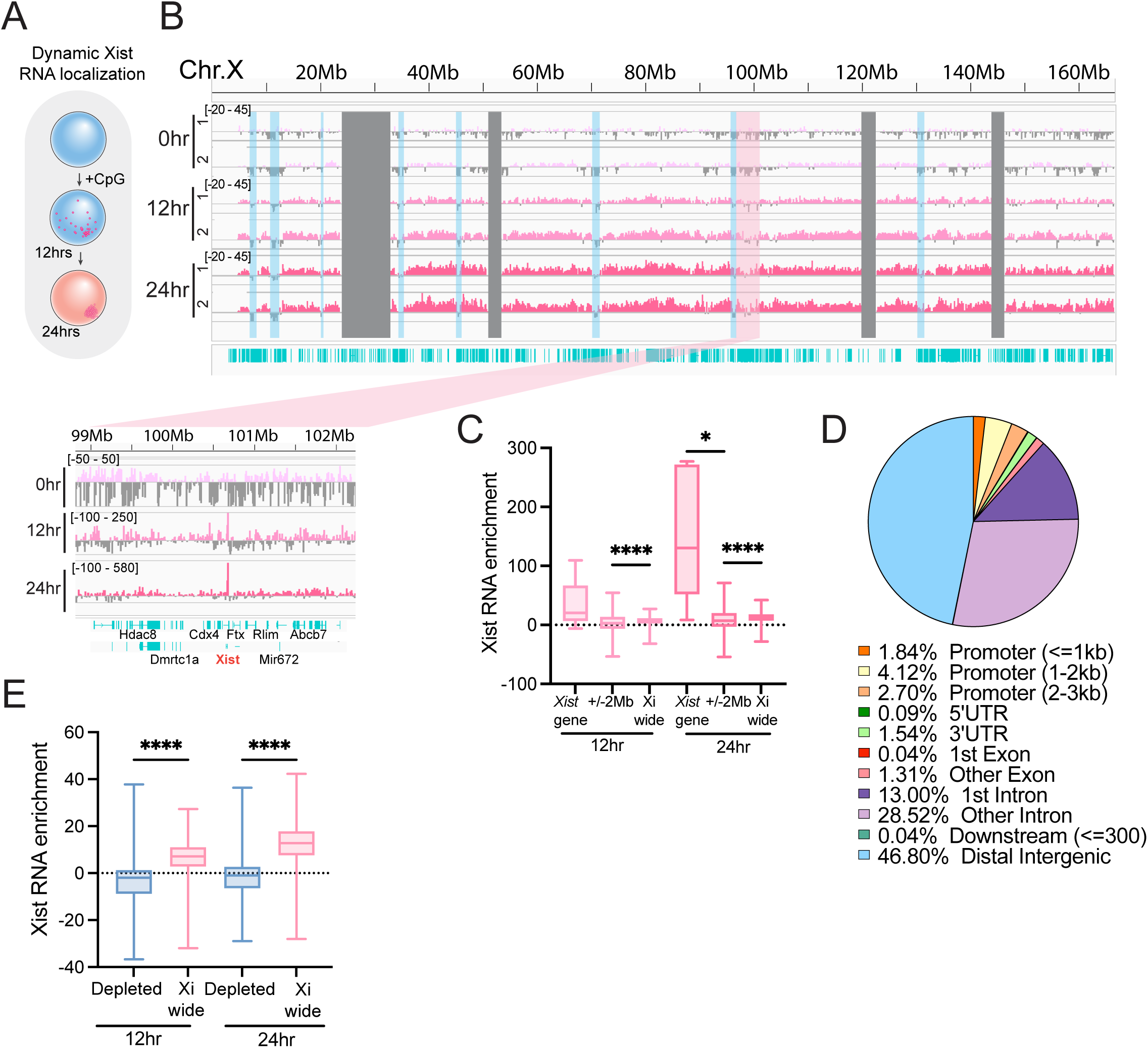
CHARTseq analysis mapping the return of Xist RNA transcripts to the Xi during B cell stimulation. **A)** Schematic of Xist RNA localization patterns during B cell stimulation observed using RNA FISH^36^. **B)** Xi-wide Xist RNA accumulation for 0, 12hr, and 24hr timepoints post-stimulation. Results from two replicate F1 animals are shown. Positive values represent the smoothed enrichment of Xist RNA over input with a scale of −20 – 40. Gray bars are un-mappable regions, blue bars highlight regions showing depleted Xist RNA. Inset image is a re-scaled and zoomed in view of Xist RNA transcripts mapping to the *Xist* promoter. **C)** Xist RNA enrichment at the *Xist* gene, a region 2Mb upstream and downstream of the *Xist* gene, and all Xi mappable regions (excluding *Xist*) using 10kb bins. Whiskers show min to max values. **D)** Annotation of mapped Xist RNA enrichment peaks from 24 hour samples (identified using MACS2 and ChIPSeeker) to genic and intergenic features across the Xi. Pie chart displays average percentages from n = 2 replicates. **E)** Comparison of Xist RNA transcript levels (‘Xist RNA enrichment’) at regions lacking detectable Xist RNA signals (blue highlighted regions in **A**) compared to all mappable regions (excluding *Xist*) on the Xi using 100kb bins. Whiskers show min to max values. All statistics were performed using a Kruskal-Wallis test, * p-value < 0.05, **** p-value < 0.0001.

### The Xi in B cells retains canonical architectural features but is more similar to the Xa than the Xi in fibroblasts

Because female B cells exhibit dynamic XCI maintenance, we next asked whether the organization of the Xi territory in naive B cells that lack Xist RNA ‘clouds’ differs from the Xi of fibroblasts and if B cell activation induced changes in Xi mega-domains or compaction. Using our F1 mouse model of skewed XCI **(Figure 1A)**, we performed allele-specific genome-wide chromosome conformation capture (Hi-C) analyses (see methods). Visual inspection of contact heatmaps at 200kb resolution indicated that while the Xa in naive and stimulated B cells exhibits the typical checkerboard pattern indicative of A/B compartments observed in autosomal chromosomes, **(Figure 3A, Supplementary Figure S3A),** the Xi in naive and stimulated B cells showed minimal attenuated compartmentalization as was previously reported for the Xi in neural progenitor cells^27^. Consistent with previous reports^27, 32^, we observed partitioning of the Xi into the two mega-domains around the *Dxz4* macrosatellite in both naive and stimulated B cells **(Figure 3A – green arrow, Figure 3B)**. However, the *Dxz4* region in stimulated B cells had a visually stronger boundary insulation for the mega-domain location compared to naive B cells (**Figure 3B)**, suggesting that cellular activation impacts boundary strength at regions across the Xi. Thus, A/B compartments exhibited significantly more fine-grained compartmentalization on the Xa than the Xi in B cells, and B cell activation did not significantly alter global compartment structure across the Xi (**Figure 3A**).

**Figure 3.**
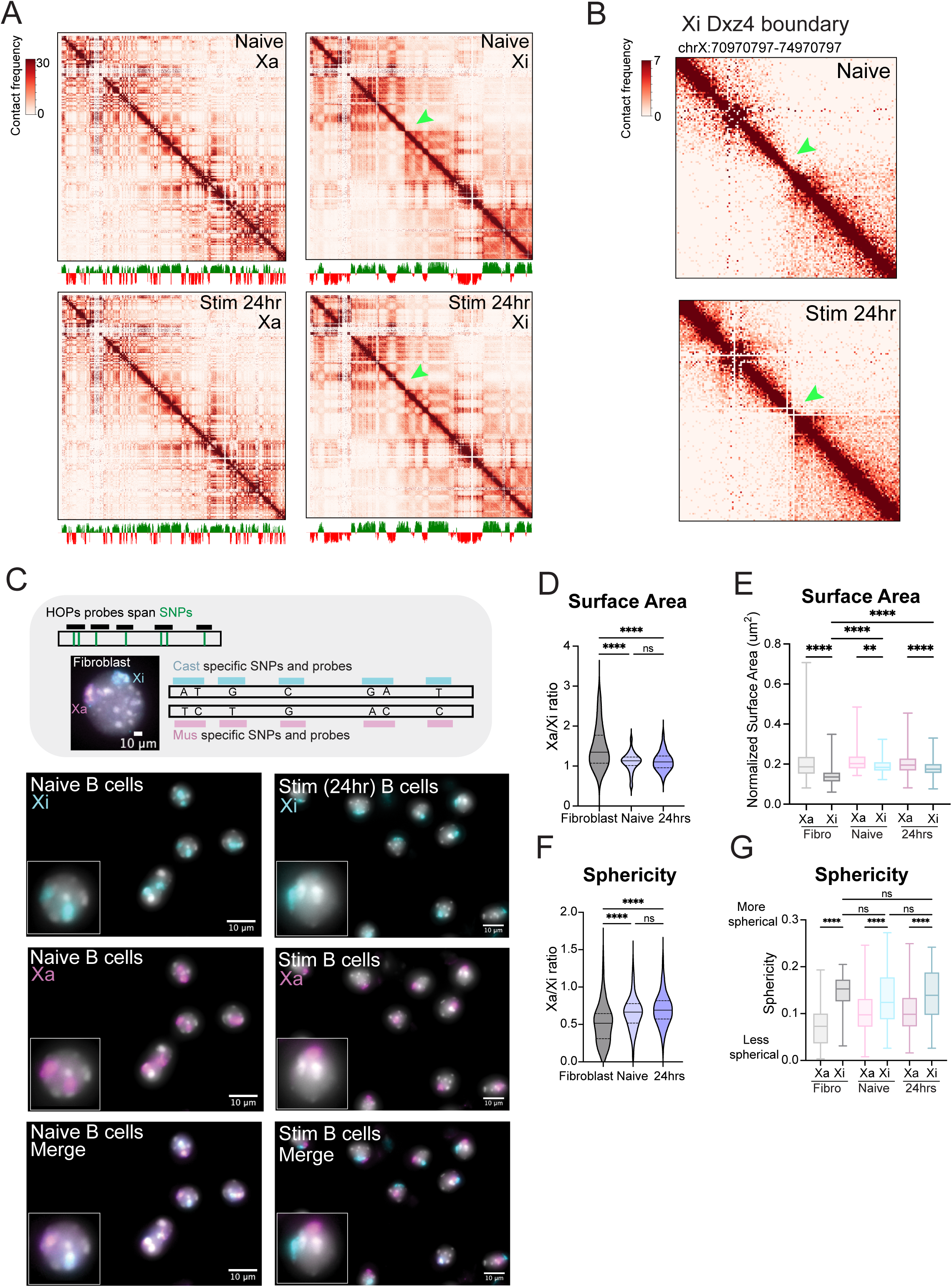
The Xi lacks A/B compartment structures, and the Xi in B cells is less compact than the Xi in primary fibroblasts. **A)** Allele-specific Hi-C heatmaps at 200kb resolution of each X chromosome from naive and *in vitro* stimulated (for 24 hours) B cells from F1 mice. Green arrow denotes *Dxz4* boundary region that separates two mega-domains on Xi. Below are A/B compartment tracks depicting A compartments in green, B in red. Scale is −5E-2 – +5E-2. **B)** Hi-C heatmaps binned at 30kb resolution showing the *Dxz4* boundary region (chrX:70970797-74970797) on the Xi in naive and stimulated (24 hours) B cells. **C)** Haplotype OligoPaints (HOPs) DNA FISH imaging for distinguishing the Xi and Xa in F1 B cells and F1 primary fibroblasts. Simplified diagram of probe design for allele-specific resolution of each X chromosome. Probes label the Xi (cyan) and the Xa (pink) in control F1 primary fibroblasts and in naive and stimulated B cells. **D)** Surface area measurements of each X chromosome territory, calculated as an Xa/Xi ratio. *** p-value <0.001. **E)** Allele-specific surface area measurements of each X chromosome territory normalized by total nuclear size. **F)** Sphericity measurements of X chromosome territories, calculated as an Xa/Xi ratio. ** p-value <0.005. **G)** Raw allele-specific measurements of sphericity for each chromosome territory. All violin plots show median with quartiles. All boxplots show median with quartiles and min to max whiskers. Statistics performed using a Kruskal-Wallis test, **p-value = 0.0089, ****p-value <0.0001. Hi-C experiments had n = 2 female mice/timepoint. Imaging experiments had n = 3 female mice/category.

As the Xi is more compact and spherical than the Xa in human diploid fibroblasts^44^, we next assessed compaction of the Xi and Xa territories in B cells using an orthogonal OligoPaints imaging method and 3D image analysis software Tools for Analysis of Nuclear Genome Organization (TANGO) to assess allele-specific chromosome territory surface area and sphericity^45–47^ in individual nuclei. We designed a library of Homologue-specific OligoPaints for X chromosomes (HOPs-X) for DNA FISH analyses^47^, in which probes contained X-linked SNPs present in either *Mus musculus* (C57Bl/6, Mus) or *Mus castaneus* (Cast) sequences **(Figure 3C – top).** Our HOPs-X probe library specifically labeled each X chromosome in primary splenic CD23+ naive and *in vitro* stimulated B cells and control primary fibroblasts isolated from F1 Mus x Cast mice **(Figure 3C)**. Although minimal off-target binding of our probes was observed, we designed a post-TANGO image analysis processing pipeline in Python that utilized the integrated density of the X chromosomes within each fluorophore to more rigorously assign the alleles as the Xa or Xi. Quantification of the surface area of the Xa and Xi chromosome territories in naive and *in vitro* stimulated B cells (24hrs) revealed an Xa/Xi ratio for surface area close to 1 for both naive (1.13 median) and stimulated (1.10 median) B cells, indicating that the surface areas of Xa and Xi were not significantly changed by B cell activation (**Figure 3D**). Normalization revealed that the surface areas of the Xi was significantly smaller than the Xa in both naive and stimulated B cells and was not affected by B cell stimulation, with the Xa median changing from 0.20 to 0.19, and the Xi median changing from 0.18 to 0.17 (**Figure 3E**). As expected, the surface area ratio (Xa/Xi) for fibroblasts was greater than 1 (1.34 median) (**Figure 3D**) and the normalized surface area of the Xi in fibroblasts was significantly lower than the Xa (median Xa = 0.19, median Xi = 0.13), but also less than the Xi surface area of B cells (**Figure 3E, Supplementary Figure S3B, S3C**).

As increased sphericity correlates with chromosome compaction^47^, we also measured the sphericity of the Xa and Xi chromosome territories in naive and stimulated B cells using HOPs-X probes. We found that the sphericity Xa/Xi ratios for naive and stimulated B cells were similar to each other (median naive = 0.66, median stimulated = 0.69), and significantly higher than the ratio in fibroblasts (median = 0.52) (**Figure 3F, Supplementary Figure S3D, S3E**). Sphericity measurements revealed that the Xi territory was consistently and significantly more spherical than the Xa territory in both B cells and fibroblasts (**Figure 3I**). B cell activation caused a slight but insignificant increase in Xi sphericity (median Xi naive = 0.12, median Xi stimulated = 0.14) but no change to the Xa (median Xa naive = 0.097, median Xa stimulated = 0.098) (**Figure 3G**, **Supplementary Figure S3D, S3E**). Together, HOPs-X measurements of the Xi and Xa territories demonstrate that the Xi is more compact and spherical than the Xa in naive and stimulated B cells and that the organization of the Xi territory in B cells is distinct from the Xi in fibroblasts.

### The Xi in naive B cells lacks TAD-like structures and B cell stimulation alters TADs and TAD boundary strength across the Xi

In NPCs and fibroblasts, the Xi lacks TADs except at regions containing XCI escape genes^27, 48^. Because we identified about 100 escape genes on the Xi **(Figure 1)** and B cell activation visually increased insulation of the mega-domain boundary (**Figure 3B)**, we asked whether the Xi exhibited TAD-like structures in B cells and whether these structures changed with B cell activation. Using our allele-specific Hi-C heatmaps binned at 30kb resolution (see methods), there was minimal folding patterns indicative of TADs across the Xi in naive B cells, even at regions of gene escape (**Figure 4A - B**). At the *Xist* region, which has the highest transcriptional level across the Xi, we observed minimal TAD-like structures in naive B cells (*Xist*; **Figure 4C**). As expected, TADs were observed across the Xa, indicating the lack of TADs on the Xi is not a general feature of chromosomes in naive B cells (**Supplementary Figure S4A - C**). We found that the Xi in stimulated B cells had visually increased TAD contacts at multiple XCI escape gene regions across the chromosome (**Figure 4A, B, green arrowheads**). To quantify stimulation-induced changes to TADs, we assessed the change in insulation score at XCI escape gene regions as well as at repressed genes on the Xi. In agreement with the visual changes in the heatmaps, we saw a significant decrease in the insulation score, indicative of increased boundary strength at XCI escape genes (**Figure 4D)**. In contrast, for transcriptionally silent genes, boundary strength (in aggregate) did not change (**Figure D**). However, there were some regions across the Xi which lacked XCI escape genes that exhibited increased TAD boundary strength (**Supplementary Figure S4D**), indicating that the gain of TADs on the Xi did not always correlate with gene expression. Thus, while the Xi in naive B cells lacks resolvable TAD structures, B cell activation induces stronger TAD boundaries on the Xi at XCI escape genes.

**Figure 4.**
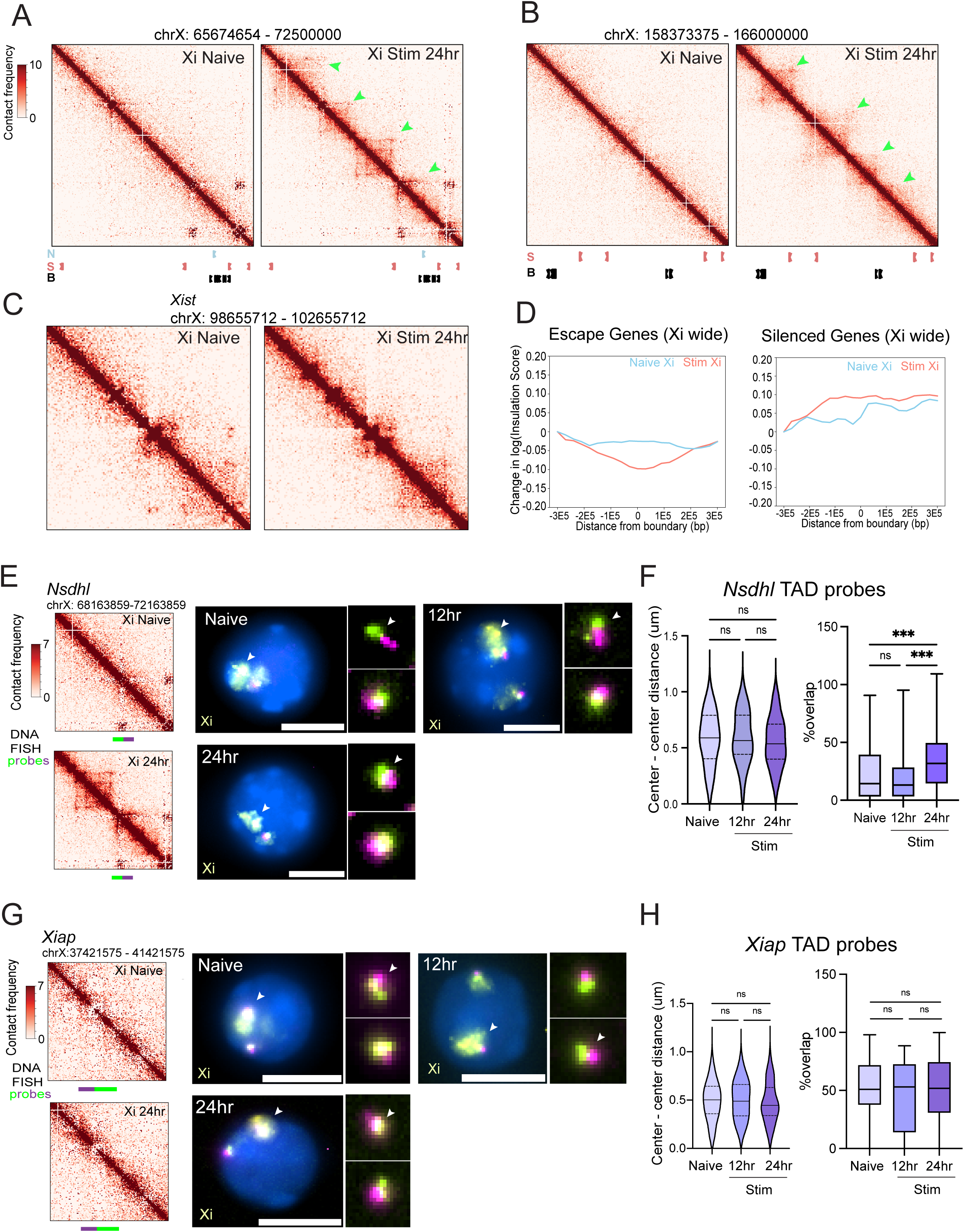
Stimulation influences TAD boundary strength on the Xi. **A)** 30kb resolution Hi-C heatmap of a region (chrX: 65674654 – 72500000) on the Xi chromosome in Naive (left) and Stim (right) B cells. Green arrows denote increased TAD interactions on the stimulated Xi. Escape genes shown below: N = naive escapee; S = stim escapee; B = escape in both. **B)** 30kb resolution Hi-C map of a egion (chrX: 158373375 – 166000000) on the inactive X chromosome in Naive (left) and Stim (right). Green arrows denote increased TAD interactions on the stimulated Xi. Escape genes shown below: S = stim escapee; B = escape in both. **C)** 30kb resolution Hi-C heatmap of a 4Mb region (chrX: 98655712 – 102655712) surrounding the *Xist* gene on the inactive X chromosome. **D)** Plot showing change in insulation score at escape genes (left) or genes subject to silencing on the Xi (right). **E)** Hi-C heatmaps of the Xi in naive and stimulated B cells centered on the *Nsdhl* region. TAD probes are shown below (green, purple). Representative images of individual nuclei using TAD probes for naive and stimulated B cells (12 hrs, 24 hrs) with Xi labeled in yellow. TADs for both alleles are shown in zoomed panels. White arrowheads denote the Xi. Scale bars are 5um. **F)** Center to center distances between *Nsdhl* TAD probes (left) and boxplot of percent overlap for the probes (right). Percent overlap is normalized to the volume of the green probe. **G)** Hi-C heatmaps of the Xi in naive and stimulated B cells centered on *Xiap* with TAD probe regions shown below (green, magenta). Right – representative images of single nuclei at naive, 12 hrs, and 24 hrs with TAD and Xi labeled in yellow. TADs for both alleles are shown in zoomed panels. White arrowheads denote the Xi-specific allele. Scale bars are 5um. **H)** Center to center distance between *Xiap* TAD probes (left) and boxplot of percent overlap (right). Percent overlap is normalized to the volume of the green probe. For imaging experiments, n=3 for 0 hr and 24 hr timepoints, n=2 for 12hr timepoint. Statistics were performed using a Kruskal-Wallis test, *** p-value < 0.0005.

To determine if the appearance of defined TAD boundaries induced by B cell activation coincided with the appearance of Xist RNA ‘clouds’ on the Xi, we used an OligoPaints DNA FISH assay to measure the spatial overlap of adjacent TAD regions, which serves as a proxy for cohesin-mediated DNA extrusion activity^49^. We designed oligos at specific Xi regions (*Nsdhl, Xiap*) at XCI escape genes that contained or lacked TAD structures in stimulated B cells. For each nucleus, we calculated both the center to center distance and the percent overlap for each signal. Loci which are closer in distance and have increased spatial overlap reflect increased extrusion activity, indicating that TAD remodeling has occurred at this region. We paired our TAD probes and HOPS-X allele-specific probes (specific to either Xa or Xi) to identify TADs on the Xi in naïve and stimulated B cells. Hi-C heatmaps for the *Nsdhl* region (ChrX:69166360-71161360) indicated that naive B cells lacked TAD-like structures on the Xi, but that TADs were present in stimulated (24 hours post-activation) B cells (**Figure 4E).** While there was a non-significant decrease in center to center distances for the TAD probes during B cell stimulation, there was a significant increase in the signal overlap at 24 hours post-stimulation (**Figure 4E, 4F**), mirroring what we saw in the Hi-C contact matrix. While the center to center differences were not significant, a trend towards decreasing probe distances at 12 hours post-stimulation is when Xist RNA transcripts begin to accumulate across the Xi (**Figure 2**), suggesting that Xist RNA re-localizes to the Xi concurrently with TAD boundary changes. For the *Xiap* region, the Hi-C heatmaps (**Figure 4G**) and TAD FISH analysis (**Figure 4H**) indicated a lack of TAD-like structures in both naive and *in vitro* stimulated B cells, as predicted. Taken together, B cell stimulation induces TAD remodeling across the Xi, coinciding with peak levels of Xist RNA transcript accumulation across the Xi.

### Xist deletion increases B cell nuclear size and impacts Xi compaction in both naive and stimulated B cells

To determine whether Xist RNA had a functional role in maintaining Xi compaction during the global de-compaction of chromosomes accompanying B cell activation ^41^, we deleted *Xist* in mature B cells by mating Xist 2lox mice^50^ to Mb1-Cre recombinase animals (with expression starting at the proB cell stage^51^). Splenic CD23+ follicular wildtype and *Xist^cKO^* B cells were activated *in vitro* using CpG and harvested at 24 hours, prior to the first cell division **(Figure 5A - top)**. As expected, we observed stimulation-induced increases in total nuclear volume and nuclear surface area for wildtype cells (**Figure 5A – bottom**). Surprisingly, we found that *Xist^cKO^* naive B cells exhibited significantly larger total nuclear volume and nuclear surface area compared to wildtype naive and stimulated B cells, and that stimulation further increased both the nuclear volume and surface area in *Xist^cKO^* cells (**Figure 5A - bottom**).

**Figure 5.**
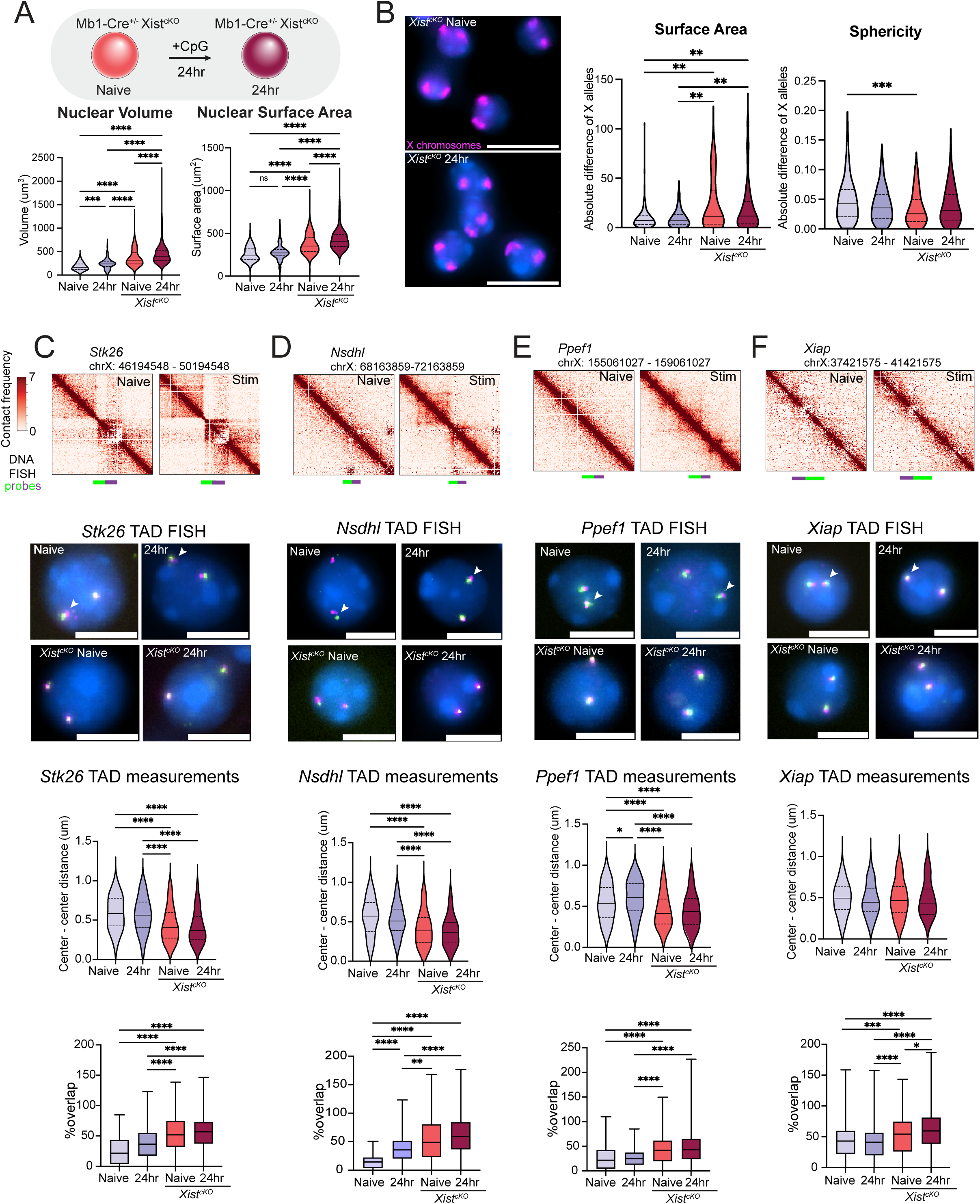
Loss of *Xist* increases nuclear size, reduces Xi compaction, and increases TAD remodeling activity in B cells. **A)** Diagram of *Xist* deletion in mature B cells using Mb1-Cre Recombinase mouse matings. Mature naive *Xist^cKO^* and *in vitro* stimulated splenic *Xist^cKO^* B cells were used for experiments. Measurements of total nuclear volume (left) and total nuclear surface area (right) for wildtype (n= 3 replicate mice) and *Xist^cKO^*cells (n= 3 replicate female mice). Statistical significance determined using Kruskal-Wallis test. *** p-value = 0.0005, **** p-value < 0.0001. **B)** Left – non allele-specific imaging of *Xist^cKO^* X chromosomes using OligoPaints. Scale bars are 10um. Right – Surface area, and sphericity measurements of the absolute difference of the X chromosomes in wild-type (purple; n=3 replicates) and *Xist^cKO^* (peach/dark purple; n = 3 replicates) samples, for both naive and stimulated (24 hrs) B cells. ** p-value < 0.005, *** p-value = 0.0008. Plots show median with quartiles. **C-F)** Hi-C heatmaps of the Xi in naive and stimulated B cells with 2-color probes (green, magenta) for measuring TAD proximity centered (+/- 2Mb) on the **C)** *Stk26 region*; **D)** Nsdhl region; **E)** Ppef1 region; **F)** Xiap region. Images show representative nuclei for each condition (n = 3 replicate mice for each genotype); TAD imaging scale bars are 5um. Measurements of center to center distances between TAD probes (above) and percent overlap between TAD probes (below). Percent overlap is normalized to the volume of the green probe within in each pair. * p-value < 0.05, ** p-value < 0.005, ***p-value = 0.0007, **** p-value < 0.0001 using a Kruskal-Wallis test. White arrowheads denote the Xi allele in F1 wild-type cells.

Based on these changes, we next asked whether *Xist* deletion would affect the compaction of the X chromosomes in either naive or stimulated B cells. Because the X chromosomes in our *Xist^cKO^* B cells cannot be distinguished using SNPs, we used DNA FISH with X-chromosome OligoPaints (**Figure 5B – left)** for single-cell detection of compaction differences that may not be detected in bulk populations. To quantify changes to Xi compaction, we calculated the absolute difference in surface area and sphericity measurements between the two X chromosomes in individual cells. A larger difference in surface area was observed between X alleles for *Xist^cKO^* naive and stimulated B cells compared to wildtype samples (**Figure 5B – middle**), suggesting that the Xi is less compact in *Xist^cKO^* B cells. B cell stimulation did not change the absolute difference of X chromosome surface areas for either *Xist^cKO^*or wildtype cells (**Figure 5B – middle**), supporting a regulatory role for *Xist* in managing Xi compaction in naive cells. Similarly, the difference in sphericity of the two X alleles was decreased in naive *Xist*^cKO^ B cells compared to wildtype naive and stimulated B cells, and B cell activation in *Xist^cKO^* cells did not further change X chromosome sphericity measurements **(Figure 5B – right**), suggesting that *Xist* deletion causes the Xi to become less spherical and less compact, and more similar to the Xa. Thus, *Xist* deletion may increase the total nuclear volume and nuclear surface area in B cells, potentially by decreasing Xi compaction and sphericity in naive B cells.

### Xist deletion increases TAD remodeling activity in naive and stimulated B cells

As Xist RNA re-localizes to the Xi concurrently with the strengthening of TAD boundaries during B cell activation (**Figure 2A, Figure 4E-H**), we at first hypothesized that Xist RNA is required to strengthen of TADs on the Xi during this process. We used our DNA FISH TAD assay to examine three regions across the X chromosome in which TAD structures emerged in stimulated B cells in Hi-C heatmaps (*Stk26, Nsdhl, Ppef1*) and one control region that lacked TADs in stimulated B cells (*Xiap*) (**Figure 5C - 5F**). We measured the center to center distances and amount of probe overlap at each region in *Xist^cKO^* naive and stimulated (24hr) B cells and compared measurements to the combined Xi and Xa results from our F1 B cells to assess TAD remodeling activity. For *Stk26, Nsdhl*, and *Ppef1* regions, we found that *Xist^cKO^* naive B cells had significantly shorter center to center distances and greater probe overlap compared to wildtype naive and stimulated B cells (**Figure 5C, 5D, 5E**), reflecting increased TAD remodeling activity. B cell activation did not change probe distances or overlap for *Xist^cKO^* cells, indicating that activation does not further increase loop extrusion/TAD remodeling seen in naïve B cells (**Figure 5C, 5D, 5E**). At the *Xiap* locus that lacks TADs in activated wildtype cells B cells, we did not observe changes in center to center distances in *Xist^cKO^* cells, but did see increased probe overlap compared to wildtype B cells (**Figure 5F**). Thus, despite the lack of localization of Xist RNA to the Xi in naïve B cells, *Xist* deletion changes the Xi structure, resulting in increased TAD boundary remodeling, likely reflecting increased frequency of TAD contacts across the Xi chromosome.

## DISCUSSION

Unlike most somatic cells, B cells utilize dynamic XCI maintenance mechanisms for dosage compensation on the Xi. During this process, Xist RNA and heterochromatic modifications, absent in naïve B cells, localize to the Xi following mitogenic stimulation prior to the first cell division. How the absence of epigenetic modifications in naive B cells and the dynamic recruitment of epigenetic modifications to the Xi following B cell activation impacts either gene expression or Xi chromatin configuration has remained unclear. Using various allele-specific approaches to identify gene expression from the Xi and assess compaction and the presence TAD-like structures, we provide the first evidence that Xi territory compaction and small scale organization across the Xi influence XCI maintenance and female biased X-linked gene expression in lymphocytes, potentially contributing to female biased autoimmune disease. In addition, our studies identify a novel role for *Xist* in regulating the territory and folding of Xi in naive and *in vitro* stimulated B cells.

As Xist RNA and heterochromatic marks are missing from the Xi in naive B cells, we speculated that the Xi would likely exhibit high levels of transcription and gene escape. Surprisingly, we find that Xi is largely dosage compensated in naive B cells, yet ∼70 X-linked genes are expressed from the Xi. There are a number of immunity-related X-linked genes that escape XCI in B cells (**Figure 1**; genes in orange) which include *Was, Il2rg, Irak1*, and *Ikbkg*, raising the intriguing possibility that increased expression of X-linked immunity genes might contribute towards loss of B cell tolerance in female-biased autoimmune diseases. Although the autoimmune-associated gene *TLR7* does escape XCI in some human B cells^39, 52^, we did not detect significant expression of *Tlr7* from the Xi in either naive or *in vitro* stimulated B cells. Despite the return of Xist RNA and heterochromatic marks to the Xi^36^, stimulated B cells have a significant number of XCI escape genes (∼93 X-linked genes), many of which also escape in naive B cells. However, more X-linked chromatin remodelers escape XCI in stimulated B cells (**Figure 1**; genes in green), including *Smc1a* and *Msl3*. As expression of these genes coincides with the appearance of TAD-like structures across the Xi, it is intriguing to consider that XCI escape of X-linked chromatin regulators contribute to genome-wide chromatin reorganization that occurs in activated B cells.

Global B cell genome reorganization occurs at 10-33 hrs post-stimulation^41^, and our work demonstrates that Xist RNA transcripts are recruited back to the Xi within this same time-frame. Using Xist CHARTseq, we confirm that Xist RNA is indeed absent across the Xi in naive B cells, as previously observed in Xist RNA FISH imaging^36, 39^, and that Xist RNA transcripts begin appearing on the Xi starting at 12 hours post stimulation (**Figure 2**). Between 12 and 24 hours, Xist RNA transcripts accumulate uniformly across the Xi (**Figure 2**), confirming our prior Xist RNA FISH imaging results^36^. Importantly, our findings are distinct from what has been described for XCI initiation during which Xist RNA first accumulates at gene-rich islands then spreads to gene poor regions^53^. As our findings indicate that dynamic XCI maintenance differs from XCI initiation, we propose that despite the lack of robust Xist RNA localization in naïve lymphocytes, epigenetic memory guides the return of Xist RNA transcripts to the Xi following B cell activation.

Using allele-specific Hi-C, we saw courser grained and attenuated compartments on the Xi in naive and *in vitro* stimulated B cells, and the existence of previously reported Xi-specific mega-domains **(Figure 3A-B)** ^27, 30, 32^. Our Hi-C experiments suggest that compartmentalization does not differ between naive and stimulated B cells, however the two mega-domains have stronger boundary insulation after CpG induced activation. Interestingly, when comparing chromosome compaction, we found that Xi territory in B cells was more similar to that of the Xa in B cells than it was to the Xi in fibroblasts (**Figure 3C-G)**, with the Xi being more compact than the Xa of B cells but less compact than the Xi of fibroblasts as assessed by both surface area and sphericity. Notably, such a difference in Xi compaction may contribute towards the increased XCI escape observed across the Xi in B cells compared to fibroblasts. Indeed, the reduced compaction of the Xi in B cells may allow for rapid gene expression changes from the Xi in response to immune stimuli.

Our work provides the first evidence that the Xi can be quickly remodeled in somatic cells, where TAD boundaries on the Xi increase in strength in response to B cell stimulation. While we did not detect TAD structures across the Xi in naive B cells in this study, it is possible that TADs could be detected across the Xi in naive B cells with higher sequencing depth. Significantly, TADs appeared in stimulated B cells which had not yet undergone the first cell division (**Figure 4**), initiated by small-scale remodeling across the Xi starting at 12 hours post-stimulation, in parallel with localization of Xist RNA transcripts across the Xi (**Figure 2**). Using immunofluorescence, we have previously shown that the heterochromatic modifications H3K27me3 and H2AK119-ubiquitin also appear on the Xi at this time^36^, which may reflect changes with TAD remodeling. In support, use of 2-color TAD specific probes for DNA FISH revealed TAD remodeling at the *Nsdhl* region and a visually stronger TAD boundary, which exhibits changes in insulation score after stimulation.

XCI escape genes are typically enriched for TAD structures, and our study found examples of XCI escape genes that contain TAD structures (*Nsdhl)* as well as those that are expressed in stimulated B cells despite lack of either TAD-like structures (*Xiap*) or evidence of TAD remodeling in stimulated B cells (**Figure 4**). In addition, our study identified repressed X-linked genes (*Ppef1)* with increased TAD interactions in Hi-C heatmaps and evidence of TAD remodeling in stimulated B cells (**Figure 5E**; wildtype cells). Thus, in stimulated B cells, TADs and TAD boundary remodeling activity on the Xi does not strictly correlate with gene expression. A recent study documented TAD formation across the Xi occurring prior to Xi reactivation during mouse iPSC reprogramming^54^, suggesting that transcription is independent of 3D chromatin organization^55, 56^. In contrast, correlations between loss of TAD structures and gene repression restriction of TADs to XCI escape regions during imprinted XCI, supports a positive correlation between Xi gene expression and TAD remodeling ^57^. In B cells with dynamic XCI maintenance, the Xi in both naive and *in vitro* stimulated B cells is dosage compensated, but only stimulated B cells have localized Xist RNA and heterochromatic marks at the Xi. However, as these stimulated cells have not yet divided, it is possible that cycle re-entry is required to observe a positive correlation between TADs and XCI escape genes. Future studies examining how abnormal overexpression of X-linked genes and loss of Xist RNA re-localization in autoimmune disease^38, 58^ impacts TAD remodeling across the Xi may provide additional insight into the relationship between transcription and chromatin organization on the Xi.

*Xist* deletion in embryonic stem cells, NPCs, and fibroblasts significantly reconfigures the Xi organization during XCI initiation and maintenance^27, 59^. However, as Xist RNA does not localize to the Xi in naive B cells, we were surprised that *Xist* deletion increased total nuclear volume and nuclear surface area in both naive and stimulated B cells (**Figure 5**). While our OligoPaint probes revealed reduced X chromosome compaction in *Xist^cKO^* B cells, it can only provide indirect evidence that the Xi chromosome is larger as we cannot distinguish X alleles in *Xist^cKO^*B cells. Therefore, we used the TAD remodeling assay to examine the impact of *Xist* deletion on TAD boundaries on the Xi. Notably, *Xist* deletion increases TAD remodeling, with greater overlap and shorter distances between TAD specific probes at all four X-loci examined, even in *Xist^cKO^* naive B cells, where Xist RNA is not localized to the Xi. As the changes to Xi organization were not further altered after *in vitro* stimulation of *Xist^cKO^* B cells, loss of *Xist* transcription may disrupt the Xi organizational structure to a level where stimulation does not have an additional impact. We envision two models for *Xist*-mediated regulation of Xi compaction and TAD formation: 1. *Xist* transcription is necessary for Xi compaction and attenuated TAD interactions across the Xi, 2. Xist RNA itself acts as a molecular scaffold for chromatin organization/remodeling complexes that bind RNAs (including Xist RNA^59^), to prevent additional binding across the Xi. Previous work demonstrated that the Xi contains fewer architectural proteins, and that Xist RNA interacts with the cohesin complex to evict cohesins from certain regions across the X chromosome during XCI initiation^59, 60^. Therefore, in the absence of Xist RNA, cohesin proteins may aberrantly accumulate across the Xi in B cells, resulting in increased looping interactions. Future work is needed to determine whether cohesin binding increases across the Xi when *Xist* is deleted in B cells and if *Xist* is necessary for maintaining Xi compaction and chromosome structure in both naive and stimulated B cells by modulating local *cis* interactions across the Xi, possibly through cohesin-mediated interactions.

## ACKNOWLEDGEMENTS

We would like to thank A. Kritz for sharing the protocol and providing invaluable input on CHART experiments; D.Beiting and PennVet CHMI for their help with sequencing; B. Gregory and X. Yu for generating the code for the allele-specific RNAseq pipeline; D.J. Emerson for advice on Hi-C analyses; L. King for help with manuscript editing; and all members from Anguera, Joyce, and Cremins labs for helpful discussions. This research was supported by an NIH R01 AI134834 (to M.C.A.), NIH 1F31GM136073-01 (to I.S.), NIGMS R35GM128903 (E.F.J.), 4D Nucleome Common Fund grants U01DA052715 U01DK127405, and R01 MH120269, 1U01DA052715, 1 DP1 MH129957 (to Phillips-Cremins).

## AUTHOR CONTRIBUTIONS

Conceptualization, I.S., M.C.A., J.E.P-C., E.F.J.; Methodology, I.S., S.C.N., M.C.A., E.F.J., J.E.P-C.; Investigation, I.S., Z.T.B.; Software, S.C.N.; Formal Analysis, I.S., H.S.R., A.L.C., R.J.B., Z.T.B.; Resources, S.C.N.; Manuscript Writing I.S. and M.C.A; Manuscript Review & Editing, I.S., M.C.A., S.C.N., R.J.B., A.L.C., E.F.J., J.E.P-C.; Funding Acquisition, I.S., M.C.A.

## Declaration of interests

The authors declare no competing interests.

## Methods

### Mice

*Xist*^fl/fl^ mice (129Sv/Jae strain) were a gift of R. Jaenisch^62^. To generate female F1 mice we used the following mating scheme: For RNAseq experiments, *Xist^fl/fl^* mice were bred to E2a-Cre (B6.FVB-Tg(EIIa-cre)C5379Lmgd/J, strain# 003724, Jackson) males to generate heterozygous *Xist^fl/+^* females. Heterozygous females were then bred with wild-type *Mus castaneus* (Cast) males to generate F1 *Xist^fl/+^* female mice. For all other allele-specific experiments, the following mating scheme was used: *Xist^fl/fl^* female mice (C57BL/6j; strain# 000664, Jackson) were mated to an ACTB-Cre male (B6N.FVB- *Tmem163^Tg(ACTB-cre)2Mrt^*/CjDswJ; strain# 019099, Jackson) to generate heterozygous *Xist^fl/+^* females. Heterozygous females were then bred with wild-type *Mus castaneus* (Cast) males to generate F1 *Xist^fl/+^* female mice. F1 *Xist^fl/+^*females from both mating schemes always inactivate the paternal WT Cast X chromosome. To generate the B cell specific knockout, *Xist*^fl/fl^ (C57BL/6j; strain# 000664, Jackson) mice were bred to an Mb1- Cre line (B6.C(Cg)-*Cd79a^tm1(cre)Reth^/*Ehobj; strain# 020505; Jackson) to generate mice heterozygous for Mb1-Cre and homozygous for *Xist^fl/fl^ (Mb1-cre^+^ Xist^fl/fl^*). All mice were maintained at the Penn Vet animal facility, and experiments were approved by the University of Pennsylvania Institutional Animal Care and Use Committee (IACUC). Euthanasia via carbon dioxide was used for animal sacrifice prior to isolations.

### B cell isolations

Follicular B cell isolations were performed as previously described using a positive selection kit^36^. Briefly, spleens from mice aged 3-6 months of age were crushed to produce single cell suspensions. Cells were incubated with Biotin tagged anti-CD23 (Clone B3B4, 101604, Biolegend), then incubated with streptavidin microbeads (130048101, Miltenyi). Cells were run through an LS column (130042401, Miltenyi) attached to a magnet. Positively selected follicular B cells were eluted from the column, and either collected immediately for experiments (0 hr/Naive timepoint) or stimulated with 1uM CpG (tlrl-1826, Invivogen) and collected at 12 hrs or 24 hrs for ‘stimulated B cell’ samples.

### Primary fibroblast isolations

Primary adult mouse fibroblasts were isolated exactly as previously published^63^. Ears from n= 3 replicate female F1 mice were removed post-mortem and cut into 3mm size pieces. Tissue was incubated in collagenase D-pronase solution [2.5mg/ml collagenase D supplemented with 250ul of 20mg/ml pronase in 4ml total] for 90min at 37C with 200rpm. Digested tissue was ground and filtered through a 70um cell strainer into complete medium [RPMI with 10% fetal calf serum, 50uM 2-mercaptoethanol, 100uM asparagine, 2mM glutamine, 1% penicillin-streptomycin]. Cells were spun for 7min at 580g at 4C and washed once with complete medium. Cells were plated with 10ml of complete medium supplemented with 10ul amphotericin B [250ug/ml stock]. Cells were cultured at 37C with 5% CO2, media was replaced on third day with fresh amphotericin B. At 70% confluency cells were split one time and sub-cultured for 2-3 days before collection and cytospinning onto slides as previously described^36^.

### Allele-specific RNAseq sequencing

B cells from n = 3 replicate F1 mus x cast female mice at either 0 hr or 24 hrs post stimulation were collected into TRIzol reagent (15596026, ThermoFisher). RNA isolations were performed according to the manufacturers protocol. Libraries were prepared with an Illumina TruSeq Stranded Total RNA LT kit (20020596, Illumina). Libraries were pooled and run on an Illumina NextSeq 500 sequencer (150bp paired-end).

To quantify gene expression from the 129S1 genome (129S1/SvlmJ, accession# ERS076385, Sanger Institute)^64^, RNA-seq reads were first aligned to the Castaneus genome (CAST/EiJ, accession# ERS076381, Sanger Institute)^64^ using STAR (v2.6.0a) with default parameters, except for the outSAMunmapped flag, which was set to Within KeepPairs to allow for unmapped reads to be extracted from alignment output. Unmapped reads were extracted using samtools and converted to Fastq format using bamToFastq. These reads were then aligned to the 129S1 genome using STAR with the outFilterMultimapNmax flag set to 1 to filter out reads that mapped to more than one loci. With the output from this second alignment, HTSeq-count (v0.10.0) was used to count allele-specific reads mapping to genes in the 129S1 genome. To quantify gene counts from the Cast genome, the same strategy was employed, except reads were first mapped to the 129S1 genome and unmapped reads were mapped to the Cast genome.

Genes that escape XCI were identified using 3 thresholds of escape, as previously described^65, 66^. Briefly, diploid gene expression was first calculated in RPKM (reads per kb of exon length, per million mapped reads), and genes were called as expressed if their diploid RPKM was > 1. For every X-linked gene that passed this threshold, haploid gene expression was calculated in SRPM (allele-specific SNP-containing exonic reads per 10 million uniquely mapped reads), and genes which had an Xi-SRPM > 2 were considered to be expressed from the Xi. Finally, a binomial model estimating the statistical confidence of escape probability was applied to the genes passing the first 2 thresholds. This model compares the proportion of Xi-specific reads to the total Xi + Xa reads and calculates a 95% confidence interval. If the 95% lower confidence limit of a gene’s escape probability was greater than 0, it was called an escapee. Genes that escaped XCI were grouped by their escape status in naive and stimulated cells, which produced 3 different categories: genes that escape in naive cells only, genes that escape in stimulated cells only, and genes that escape in both naiveand stimulated cells.

Reads were graphed using Prism v9.3.1, statistical significance was determined using one-way ANOVA with Tukey’s multiple comparison test. The R package chromoMap was used to generate chromosome maps displaying aggregate RPKMs^61^. Venn diagraph of escape genes was generated using VennDiagram package in R. Heatmaps were generated using RPM values of genes as input to the gplots function heatmap.2 in R with data scaling set to “row”. GO analysis was performed using Metascape^67^.

### Capture hybridization of RNA targets (CHART)

CHART protocol was performed as previously described^15, 30^. Splenic follicular B cells were isolated at 0 hr, 12 hrs, and 24 hrs post CpG stimulation (n=2 mice per timepoint, 25 million cells/replicate). Cells were crosslinked in 1% formaldehyde for 10min at room temperature. Cells were incubated in sucrose buffer [0.3M sucrose, 1% Triton X-100, 10mM HEPES, 100mM KOAc, 0.1mM EGTA, 0.5mM Spermidine, 0.15mM Spermine, 1x complete EDTA-PIC, 10U/ml SUPERasIn (AM2696, Thermo Scientific)] and nuclei isolated through 20 passes in a dounce homogenizer with a tight pestle. Nuclei were collected through centrifugation in glycerol cushion [25% glycerol, 10mM HEPES, 1mM EDTA, 0.1mM EGTA, 100mM KOAc, 0.5mM Spermidine, 0.15mM Spermine, 1x complete EDTA-PIC, 1mM DTT, 5U/ml SUPERasIn]. Nuclei were crosslinked again in 3% formaldehyde for 30min at room temperature. Crosslinked nuclei were incubated for 10min at 4C in nuclear extraction buffer [50mM HEPES, 250mM NaCl, 0.1mM EGTA, 0.5% N-lauroylsarcosine, 0.1% sodium deoxycholate, 5mM DTT, 10U/ml SUPERasIn]. Nuclei were spun down and resuspended in sonication buffer [50mM HEPES, 75mM NaCl, 0.1mM EGTA, 0.5% N-lauroylsarcosine, 0.1% sodium deoxycholate, 0.1% SDS, 5mM DTT, 10U/ml SUPERasIn], and sonicated in a Covaris S220 sonicator with the following conditions: PIP – 140W; Duty factor – 10%; Cycles – 200; for a total of 8mins at 4C [Of note, the 0 hr samples required a longer sonication time of 10mins]. Sonicated lysates were pre-cleared with MyOne Streptavidin C1 beads (65001, Thermo Scientific) for 1hr at room temperature in 2X hybridization buffer [50mM Tris-HCl pH 7.0, 750mM NaCl, 1% SDS, 1mM EDTA, 15% formamide, 1mM DTT, 1mM PMSF, 1X PIC, and 100U/mL SUPERaseIN]. Input [1%] was removed and frozen at −80C. Pre-cleared lysates were then incubated with a pool of 10 biotinylated Xist oligos^15^ (see Supplementary Table 3 for sequences) at a final concentration of 36pmol. Samples were incubated for 4hr at 37C. Samples were washed once in 1X hyb buffer [33% sonication buffer, 67% 2X hybridization buffer]; five times with 2% SDS wash buffer [10mM HEPES, 150mM NaCl, 2% SDS, 2mM EDTA, 2mM EGTA, 1mM DTT]; and two times with 0.5% NP40 wash buffer [10mM HEPES, 150mM NaCl, 0.5% NP40, 3mM MgCl2, 10mM DTT] at 37C. Beads were resuspended with 200ul 0.5% NP40 buffer and DNA was eluted twice from beads using 20ul RNase H [5U/ul] at 37C for 30min each time. Input and eluted DNA were treated with 10ul RNase A [20mg/ml] at 37C for one hour, followed by addition of 10ul Proteinase K [20mg/ml] and incubated at 55C for 1hr. After, 12ul 5M NaCl was added and samples were reverse crosslinked at 65C overnight. DNA was isolated following a standard phenol-chloroform extraction.

### CHART sequencing and analysis

Library preparation was preformed using the NEB Next Ultra II library prep kit. For starting material, the same concentration of DNA was used between each samples input and IP values as measured by Qubit. The samples were sequenced on a NextSeq 2000 with 2×150bp read length.

For analysis, FastQ files were trimmed using Trim Galore! (https://www.bioinformatics.babraham.ac.uk/projects/trim_galore/) and Cutadapt to remove adapter sequences and low quality reads and checked using FastQC. Trimmed FastQ files were aligned to the mm9 genome using Bowtie2 v2.3.4.1^68^. Aligned output files were sorted using samtools^69^ and filtered using sambamba^70^ to remove duplicates, unmapped reads, reads mapping to mitochondrial DNA, and improper pairs. Blacklist filtering^71^ was performed using bedtools intersect^72^. Sorted and filtered BAM files were analyzed in R using SPP^73^. To generate enrichment files the function get.smoothed.tag.density was used with smoothing using 1Mb windows every 500bp with input files as controls. To control for sequencing depth, files were scaled using total positive read density on chromosome 4^15^. Enrichment files were converted to BigWig format and visualized using IGV^74^. For determining Xist RNA enrichment at specific genomic regions, mappable regions were binned using bedtools makewindows at either a 10kb or 100kb window size (specified in figure legends). The *Xist* locus was excluded from the total X comparison to prevent skewing of the enrichment values. deepTools2 multiBigwigSummary^75^ was then used to extract enrichment values using the BED file outputs from bedtools. To determine genomic annotation of enrichment, peak calling was performed using the MACS2^76^ function callpeak with paired-end BAM flag (-f BAMPE), keep duplicates set to auto, a q-value cutoff of 0.1, and with scaling set to small. Output peak file was used with the Bioconductor package ChIPseeker^77^ to determine genomic annotations, with pie chart generated using Prism v9.3.1. UCSC Table Browser^78^ was used to find genes located in Xist RNA depleted regions. Genes were considered expressed if the 129S1/Xa RPKM values were >0 in at least two samples. Graphs were created using Prism v9.3.1, Kruskal-Wallis statistical tests were performed to determine significance.

### Hi-C sequencing

Hi-C was performed using the Arima Hi-C+ kit (Arima Genomics, San Diego, CA, USA) following their standard protocol. 2 million cells per replicate (n = 2 mice for each timepoint) were crosslinked with 2% formaldehyde for 10 mins at RT before proceeding with Hi-C. For library preparation, the KAPA HyperPrep kit (07962312001, Roche) was used with a modified protocol provided by Arima Genomics. Libraries were checked by Tapestation analysis, pooled, and run for two rounds on a NextSeq 500 with a 2×150bp read length. Two replicates were pooled to generate 400 million reads/timepoint.

### Allele-specific Hi-C analysis

We performed allele-specific Hi-C analysis using Hi-C-Pro v2.8.9 according to the allele-specific analysis section of the Hi-C-Pro manual (https://nservant.github.io/Hi-C-Pro/AS.html). Briefly, we generated a masked mm9 genome, in which each location of strain-specific SNPs differentiating CAST and C57Bl6 strains is N-masked. First, the NCBI37/mm9 genome was downloaded from the UCSC Genome Browser (https://hgdownload.soe.ucsc.edu/downloads.html#mouse). We then downloaded a database of strain-specific SNPs for a variety of mouse strains from the Sanger Institute Mouse Genomes Project (https://www.sanger.ac.uk/data/mouse-genomes-project/)^64^. Using this information, we generated a vcf file by running the extract_snps script on Hi-C-Pro that contained SNPs that differ between CAST and C57bl6 mouse genome. We masked the reference mm9 genome at all loci of SNPs specified in the vcf file by running the bedtools maskfasta command (https://bedtools.readthedocs.io/en/latest/content/tools/maskfasta.html). Finally, we built a Bowtie index of the masked genome using the bowtie2-build indexer (http://bowtie-bio.sourceforge.net/bowtie2/manual.shtml#the-bowtie2-build-indexer).

We aligned paired-end reads from Hi-C fastq files to our masked mm9 genome using bowtie2 (global parameters: --very-sensitive -L 30 --score-min L,-0.6,-0.2 --end-to-end – reorder; local parameters: --very-sensitive -L 20 --score-min L,-0.6,-0.2 --end-to-end – reorder). We filtered unmapped reads, non-uniquely mapped reads, and PCR duplicates (Supplementary Table 4).

We assembled raw cis contact matrices for each chromosome at two different time points for each allele at 30kb and 200kb resolution. X chromosome alleles were assigned such that Xa represented the C57bl6-specific allele and Xi represented the CAST-specific allele. Replicates for each condition were merged. We balanced the merged contact matrices using Knight-Ruiz balancing as previously described^79, 80^. For each balanced matrix, we performed simple scalar normalization, where a simple scalar size factor for each pixel was calculated based on the genomic distance between pairs of bins^81^.

### Domain calling

We used our previously published domain caller 3DNetMod^80^ (https://bitbucket.org/creminslab/cremins_lab_tadsubtad_calling_pipeline_11_6_2021/src/master/) to identify TADs and subTADs on normalized, balanced Hi-C matrices binned at 30 kb resolution. We log transformed counts and chunked the data into both 6 Mb regions with 4 Mb overlap between adjacent regions and 3 Mb regions with 2 Mb overlap^80, 82, 83^. As previously described, we filtered sparse regions^83^. We considered regions to be sparse if a chunked region contained zero counts for 1/3 of all pixels on the diagonal or if it contained consecutive zeros for more than a 1500kb distance. We identified high-confidence domains by using gamma steps of 0.01 to perform a ‘gamma plateau sweep’ which compares distribution of communities of domains identified genome-wide^83^. Plateaus were identified as consecutive gamma steps resulting in the same number of communities (mean per 20 partitions). We required a minimum plateau size of 16 and 8 for 6 Mb and 3 Mb chunked regions, respectively. We then merged the 6 Mb and 3 Mb chunked regions and filtered out domains smaller than or equal to 150 kb. We created a final set of unique boundary locations by merging redundant domains and colocalizing boundary locations. Domains were considered redundant if two domains were within +/- 60 kb on both boundary edges. Boundary locations were colocalized to share a single consistent boundary if the gap between adjacent boundaries was less than 7.5% of the domain size of each of the boundaries or if boundaries were located within 60 kb of each other.

### Compartment calling

To plot A/B compartment tracks chromosome-wide, we performed eigenvector decomposition on 200 kb resolution, balanced Hi-C matrices, as was previously described^84–87^. In short, matrices were normalized using a global expected distance dependence mean counts value. Low coverage rows and columns were filtered, and we transformed off-diagonal counts to obtain a z-score, which was used to generate a Pearson correlation matrix. Finally, we performed eigenvector decomposition on the resultant matrix.

### Boundary strength with insulation score

As previously described^84^, we calculated insulation scores chromosome-wide by applying a 300 kb square summation window with 30 kb offset in 30 kb resolution, balanced Hi-C data^88^. Insufficient counts occurring at the beginning and end of the chromosome were discarded. We constructed aggregate plots of mean insulation scores centered on genes (+/- 300 kb around the center of each gene) for escape genes and silenced genes chromosome-wide in Xi.

### Data availability

All sequencing data generated in this study has been deposited to the NCBI GEO database. Access data using the following accession numbers: GSE215848 [Hi-C, CHART] and GSE208393 [RNAseq].

### Allele-specific HOPs probe libraries

Probes were mined for the X chromosome (mm10 genome build) using the OligoMiner design pipeline^89^, with the -l and -L parameters set to 42 for 42-mer oligos and the -O parameter added for overlapping oligos. The resulting set of oligos were then modified to include SNPS for the Cast/EiJ or C57BL/6NJ mouse strains, downloaded from the Mouse Genomes Project (https://www.sanger.ac.uk/data/mouse-genomes-project/). Strain-specific Oligopaints were selected using a similar workflow to the HOPs pipeline^90^. Specifically, oligos were selected based on containing at least one differential SNP in the inner 32 nucleotides of each 42-mer oligo. These oligos were purchased from CustomArray/Genscript, and probe sets were produced as described previously^46^.

### DNA FISH with OligoPaints

Splenic B cells (0 hr and 24 hr post CpG stimulation) from n = 3 replicate female mice were cytospun onto slides and processed as previously described^36^. Slides were briefly incubated in room temperature SSCT+formamide [2X SSC, 0.1% Tween-20, 50% formamide], then pre-hybridized in SSCT+formamide for 1hr at 37C. Primary probe mix [50% formamide, 1X Dextran Sulfate Mix [10% dextran sulfate, 4% PVSA, 2X SSC, 0.1% Tween-20], 10ug RNase A, 5.6mM dNTPS, 50pmol per Oligopaint probe was added to slides, sealed with rubber cement, and denatured for 30min at 80C. Slides were then hybridized overnight at 37C in a humidified chamber. Next day, slides were washed for 15min in 2X SSCT [2X SSC, 0.1% Tween-20] at 60C, 10min in 2X SSCT at room temperature, then 10min in 0.2X SSC at room temperature. Secondary probe mix [10% formamide, 1X Dextran Sulfate Mix, 10pmol per secondary probe] was added to slides, sealed with rubber cement, and incubated in a humidified chamber for at least 2hr at room temperature. Slides were washed for 5min in 2X SSCT at 60C, 5min in 2X SSCT at room temperature, then 5min in 0.2X SSC at room temperature. Slides were mounted with Vectashield and imaged on a Nikon Eclipse microscope with Z-stacks set to a 0.2um step size.

### Image analysis

All images were analyzed using TANGO^91^. The following settings were used for allele-specific images and whole X imaging in *Xist* conditional knockout cells: Nuclei – pre-filter: Fast Filters 3D; Segmentation: Hysteresis Segmenter; Post-filters: Size and Edge Filter, Morphological Filters 3D (Fill Holes 2D, Binary Close). Alleles – pre-filter: None; Segmentation: Hysteresis Segmenter; Post-filters: Size and Edge filter, Erase Spots. The following settings were used for TAD imaging: Nuclei – pre-filter: Fast Filters 3D; Segmentation: Hysteresis Segmenter; Post-filters: Size and Edge Filter, Morphological Filters 3D (Fill Holes 2D, Binary Close). TADs – pre-filter: Fast Filters 3D, Gaussian Smooth; Segmentation: Hysteresis Segmenter; Post-filters: Size and Edge filter, Erase Spots.

For allele-specific X chromosome imaging, TANGO-generated raw allele-specific data files were further processed using a custom python script which utilizes the integrated density to select the true allele for each genome. Cells with overlapping chromosomes were excluded from analyses. For surface area measurements, each allele was normalized to its respective nuclear size. To generate surface area and sphericity ratios, the active X value was divided by the inactive X value within each individual cell then graphed. For proportion plots, the allele of choice (Xn) was divided by the sum of values for both alleles for each particular measurement (Xn/Xi+Xa). Graphs were created using the ecdf function in R. Whole X imaging in *Xist* conditional knockout cells were analyzed post segmentation by taking the absolute difference of alleles in each nucleus for each measurement. The same analysis was performed on the allele-specific data for comparison.

For all TAD imaging, alleles were filtered out if their center – to – center distance was greater than 1um (removed trans measurements between alleles), and only cells which contained two distinct alleles (two objects for each probe) were used for further analysis. For allele-specific TAD imaging, whole X allele-specific probes were used in conjunction with TAD probes, but only the Xi allele was labeled with a secondary probe. The Xi- specific TADs were manually selected by proximity to the labeled Xi allele. Distances were measured using the center-to-center distance between each TAD within each allele. Overlap values were normalized to the volume of one probe from each set (indicated in figure legends).

Unless specified above, all graphs were generated using Prism v9.3.1. Significance was determined by Kruskal-Wallis statistical tests performed in Prism v9.3.1.

### Data availability

All sequencing data generated in this study has been deposited to the NCBI GEO database. Access data using the following accession numbers: GSE215848 [Hi-C, CHART] and GSE208393 [RNAseq].

---

**Figure S1.**
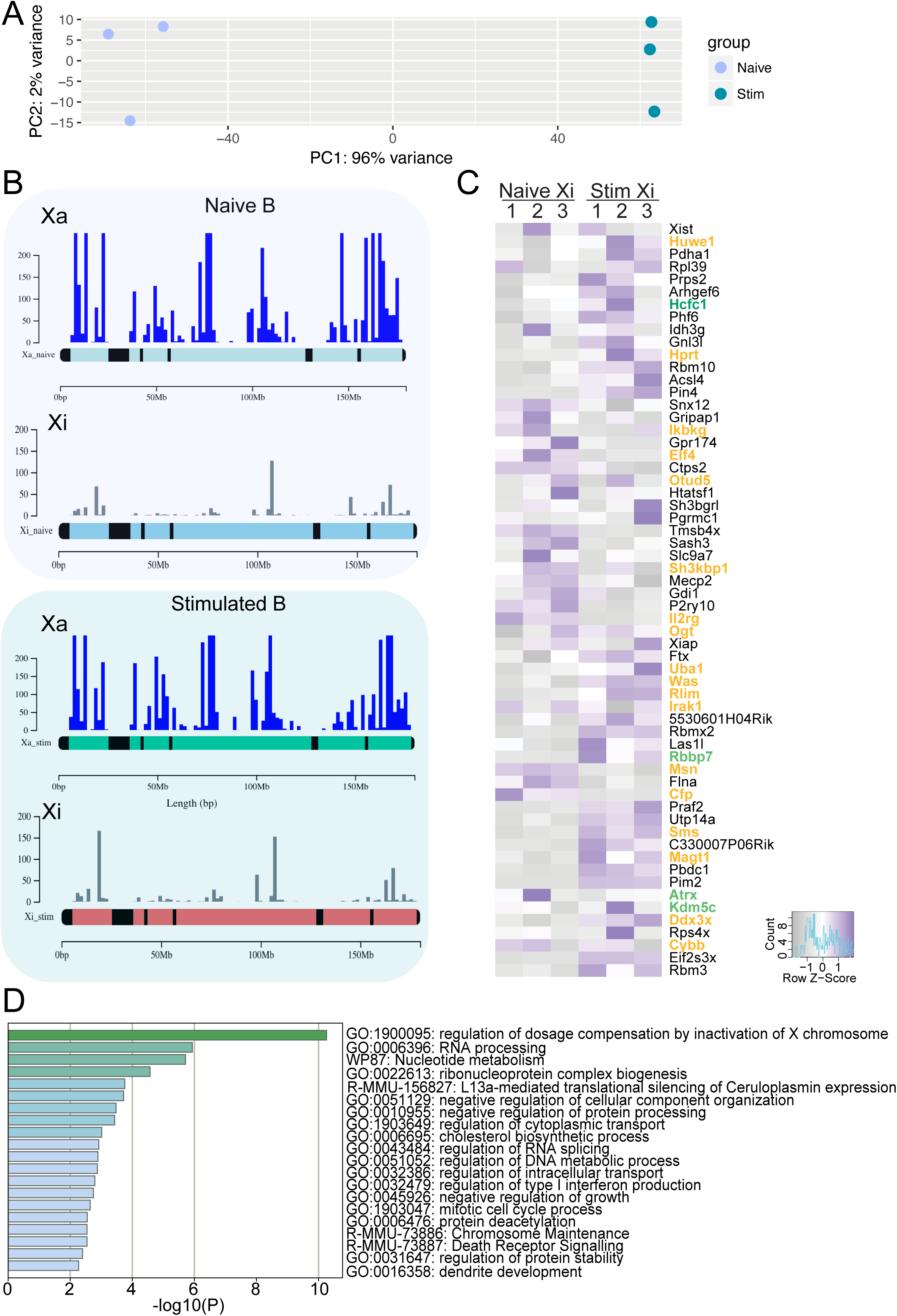
XCI escape genes in naive and stimulated B cells. **A)** PCA plot of naive (purple) and 24 hour stimulated (teal) B cells. **B)** ChromoMaps comparing aggregate RPKM expression of the Xa and the Xi in naive (top) and stimulated (bottom) B cells. The Xa is scaled to match the Xi. **C)** Heatmap showing z-scores of the Xi-specific expression of the 60 XCI escape genes expressed in both naive and stimulated B cells. Genes in orange have immunity-related functions; genes in green have roles in chromatin organization. **D)** Metascape analysis of all 104 XCI escape genes in B cells.

**Figure S2.**
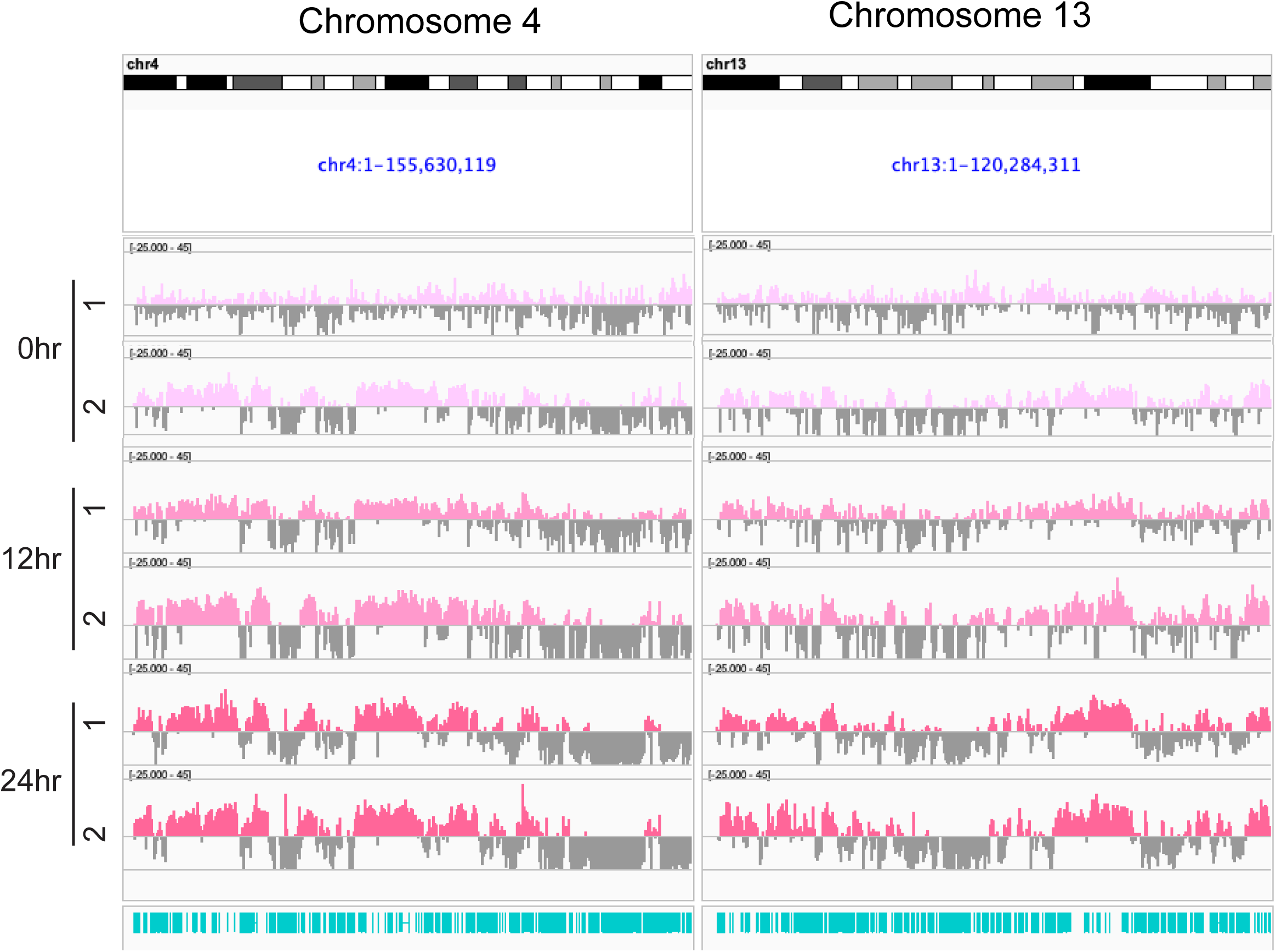
Xist RNA CHARTseq showing background of Xist RNA transcripts across two autosomal chromosomes. Xist RNA CHARTseq results for n=2 naive and stimulated B cells at 12 and 24 hour timepoints on chromosomes 4 and 13. Positive values represent the smoothed enrichment of Xist RNA over input with a scale of −20 – 40; gray bars are un-mappable regions. The Xist RNA transcripts mapping to chromosomes 4 and 13 are background, and this level of enrichment is similar to the X chromosome in naive B cell samples in Figure 2.

**Figure S3.**
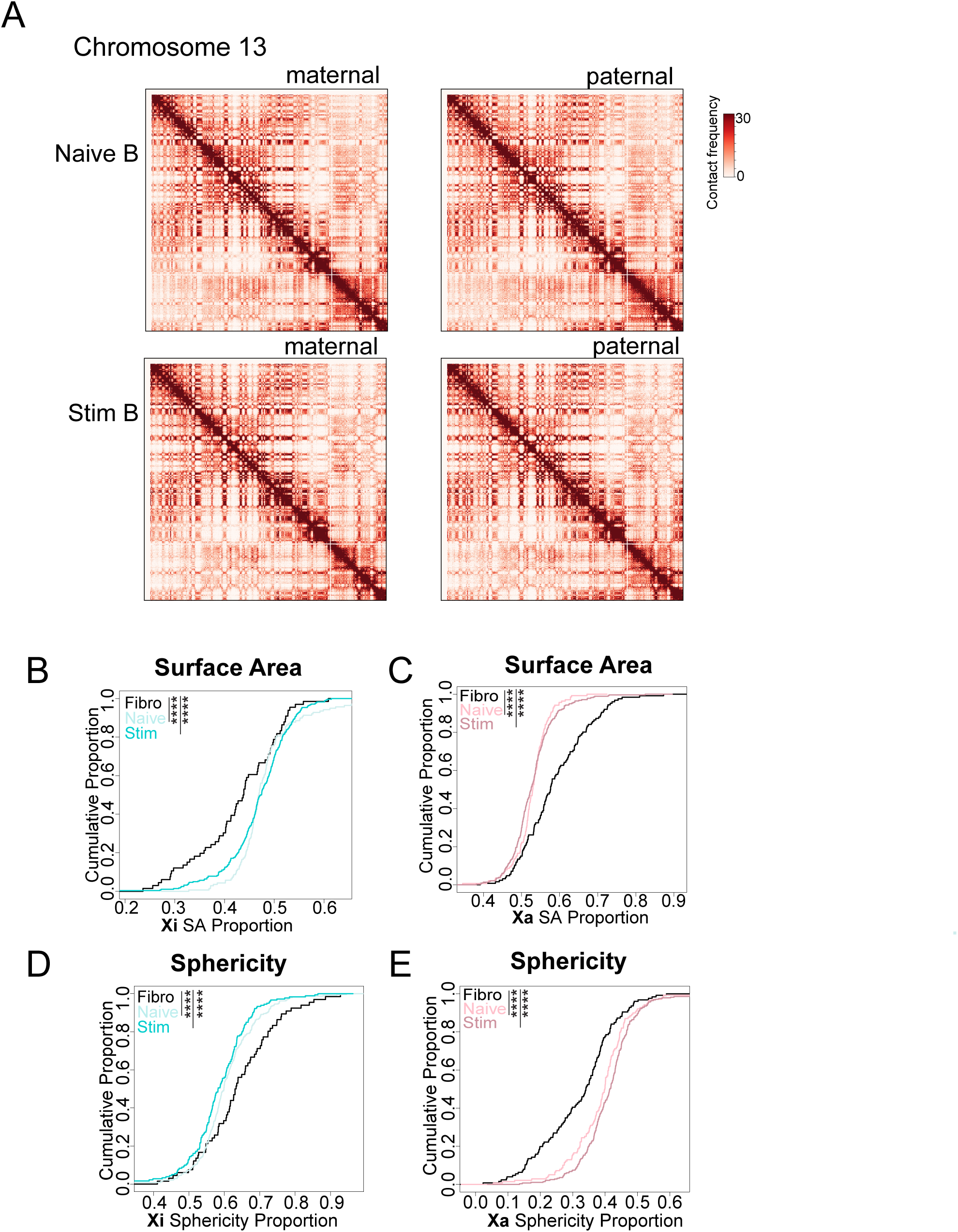
Allele-specific Hi-C maps for chromosome 13 and cumulative distributions of imaging measurements for Xi and Xa. **A)** 200kb resolution Hi-C heatmaps of Chromosome 13 displaying both maternal (Mus) and paternal (Cast) alleles for naive and 24 hour stimulated B cells. **B)** Cumulative proportion plots for surface area of the Xi relative to total X chromosome surface area in fibroblasts (black), naive B cells (light cyan), and stimulated B cells (dark cyan). **C)** Cumulative proportion plots for surface area of the Xa to total X chromosome surface area in fibroblasts (black), naive B cells (light coral), and stimulated B cells (dark coral). **D)** Cumulative proportion plots for sphericity of the Xi to total X chromosome in fibroblasts (black), naive B cells (light cyan), and stimulated B cells (dark cyan). **E)** Cumulative proportion plots for sphericity of the Xa relative to total X chromosome in fibroblasts (black), naive B cells (light coral), and stimulated B cells (dark coral). Significance was determined by Kruskal-Wallis test, **** p-value <0.0001.

**Figure S4.**
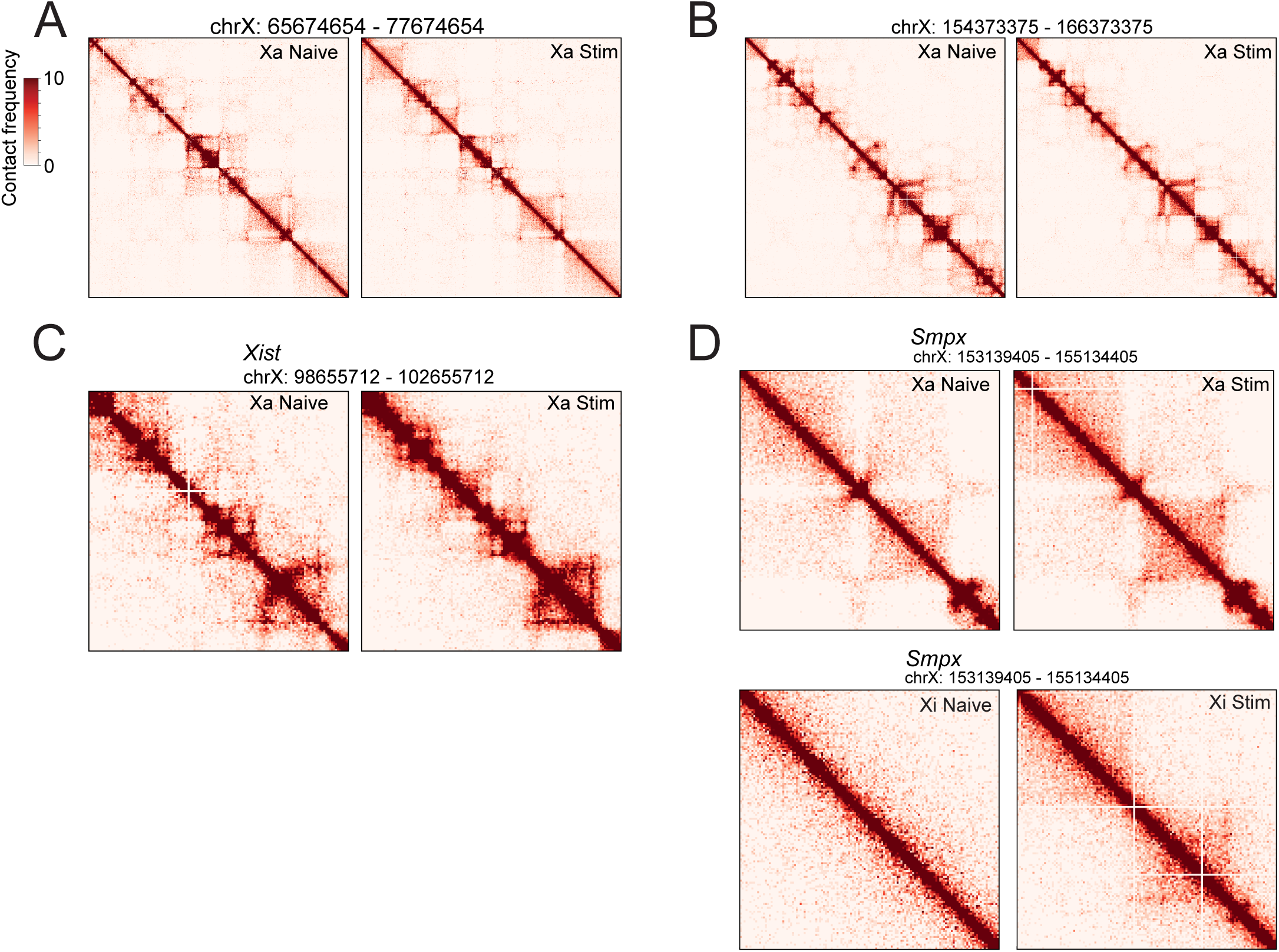
Hi-C maps showing TAD structures on the Xa and plots of gene enrichment at TADs on the Xi. **A)** 30kb Hi-C heatmap of a 12Mb region (chrX: 65674654 – 77674654) on the Xa in naive (left) and 24 hrs stimulated (stim) B cells. **B)** 30kb Hi-C map of a 12Mb region (chrX: 154373375 – 166373375) on the Xa in naive (left) and stimulated B cells (right). **C)** 30kb Hi-C heatmap of a 4Mb region (chrX: 98655712 – 102655712) encompassing the *Xist* gene on the Xa. There is a region of *Xist* deleted on the Xa, and therefore this map may not represent the true structure of the wild-type *Xist* locus on the Xa. **D)** 30kb Hi-C maps of a 4Mb region (chrX: 152136905 – 156136905) centered on the transcriptionally silent gene region of *Smpx (*+/- 2Mb) for both the Xa (top) and the Xi (bottom) in naive and stimulated B cells.

## Notes

### Competing Interest Statement

The authors have declared no competing interest.

